# Joint probabilistic modeling of pseudobulk and single-cell transcriptomics enables accurate estimation of cell type composition

**DOI:** 10.1101/2025.05.28.656123

**Authors:** Simon Grouard, Khalil Ouardini, Yann Rodriguez, Jean-Philippe Vert, Almudena Espin-Perez

## Abstract

Bulk RNA sequencing provides an averaged gene expression profile of the numerous cells in a tissue sample, obscuring critical information about cellular heterogeneity. Computational deconvolution methods can estimate cell type proportions in bulk samples, but current approaches can lack precision in key scenarios due to simplistic statistical assumptions, limited modeling of cell-type heterogeneity and poor handling of rare populations. We present MixupVI, a deep generative model that learns representations of single-cell transcriptomic data and introduces a mixup-based regularization to enable reference-free deconvolution of bulk samples. Our method creates a latent representation with an additive property, where the representation of a pseudobulk sample corresponds to the weighted sum of its constituent cell types. We demonstrate how MixupVI enables accurate estimation of cell type proportions through benchmarking on pseudobulks simulated from a large immune single-cell atlas. To support reproducibility and foster progress in the field, we also release PyDeconv, a Python library that implements multiple state-of-the-art deconvolution algorithms and provides a comprehensive benchmark on simulated pseudobulk datasets.

## 1. Introduction

The cellular composition of tissues plays a critical role in their function, and alterations in this composition are associated with various diseases including cancers (Li, 2017; Rooney, 2015; Gentles & Newman, 2015; Mahmoud, 2011; Li, 2016), cardiovascular diseases (Parker, 2020), and neurodegenerative disorders (Lee, 2012; Poduri, 2012; Erickson, 2010; Rivière, 2012). Bulk RNA sequencing (RNA-seq) has been widely used to profile gene expression in tissue samples (Cancer Genome Atlas Research Network, 2013; Greenwood, 2020; GTEx Consortium, 2020), but this technology measures the average expression of all cells in a sample, masking the heterogeneity of different cell types. In contrast, single-cell RNA sequencing (scRNA-seq) provides gene expression profiles of individual cells, allowing direct characterization of cellular heterogeneity. However, scRNA-seq is more expensive, technically challenging, and often limited in the number of samples that can be processed compared to bulk RNA-seq.

Computational deconvolution methods aim to bridge this gap by estimating cell type proportions in bulk RNA-seq samples using reference profiles derived from single-cell data (Cobos, 2020). Traditional approaches rely on linear regression models that express bulk samples as linear combinations of these reference profiles, often referred to as a signature matrix (Erdmann-Pham, 2021; Dong, 2021; Tsoucas, 2019; Newman, 2015). While these approaches have shown promise, they face several limitations. In particular, they often fail to account for the complex differences between single-cell and bulk RNA-seq data, biological variations within cell types or due to simplistic statistical assumptions (e.g modeling transcriptomic data with additive Gaussian noise).

Deep generative models have recently emerged as powerful tools for analyzing single-cell data (Donno, 2023; Cui & Wang, 2024), enabling the learning of low-dimensional representations that capture biological variation while accounting for technical factors (Xu, 2021). Models such as single-cell Variational Inference (scVI) have demonstrated consistent performance in tasks such as clustering, batch correction, and imputation of single-cell data (Lopez, 2018). However, reconciling the expressive nature of these non-linear deep generative models with the linearity constraints imposed by the deconvolution problem remains a challenge.

To address this gap, we introduce MixupVI, a deep generative model for bulk RNA-seq deconvolution that leverages single-cell representations. Our approach constrains the latent space of a variational autoencoder to enforce an additive property, where the representation of a bulk sample can be approximately expressed as a weighted sum of cell-type specific latent representations. This enables accurate estimation of cell type proportions in pseudobulk samples while maintaining the benefits of deep generative modeling.

### 1.1. Conventional assumptions in deconvolution approaches

Bulk transcriptomics deconvolution aims to estimate the proportions of distinct cell types within a heterogeneous tissue sample based on its aggregated gene expression profiles. Traditional deconvolution methods build on Least Squares approaches. These techniques rely on the critical assumption of linearity, which states that the expression level of each gene in a bulk sample is a linear combination of the expression levels from the constituent cell types, weighted by their proportions. Formally, let *G* denote the number of genes and *C* the number of cell types in the deconvolution setting. The gene expression profile of a bulk sample *b* can be modeled as:

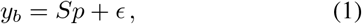

where:

- *y*_*b*_ ∈ ℕ^*G*^ is the observed bulk gene expression vector,
- *S* ∈ ℝ^*G×C*^ is the cell-type specific reference (signature) matrix,
- *p* ∈ ℝ^*C*^ is the vector of cell type proportions (with ∑_*c*_ *p*_*c*_ = 1),
- *ϵ* ∼ 𝒩 (0, *σ*^2^*I*_*G*_) is Gaussian noise.

The assumption of linearity holds reasonably well in practice, especially when enough cells are present in a sample and the deconvolution granularity and observed proportions are appropriate. However, this assumption can break down in key scenarios, leading to systematic biases in cell type proportion estimates:

- Log-transformation is commonly used in transcriptomics to stabilize variance and approximate Gaussianity to rely on Least Squares approaches. However, it distorts the linear relationships between mixed cell populations in bulk data. This results in curved trajectories in PCA space (Supplementary Figure S1), leading to systematic errors in proportion estimates - especially for intermediate mixtures. As a result, some deconvolution methods advice to keep the data in linear space (Zhong & Liu, 2011; Newman, 2015). The data thus no longer follows a Gaussian distribution and instead typically exhibits over-dispersed negative binomial behavior with prominent zero-inflation.
- Low abundance. When a cell type is barely present within a sample, its gene expression signal may be too weak or noisy to resemble the average profile captured in the reference signature matrix.
- Low granularity. Broad cell type definitions - such as grouping all lymphoid cells together - fail to capture intra-lineage heterogeneity. While CD4^+^ T cells, CD8^+^ T cells and Natural Killer cells originate from the common lymphoid progenitor, they exhibit distinct gene expression profiles and unique markers (Hidalgo, 2008). Overlooking these differences can increase variance and reduce deconvolution accuracy (Supplementary Figure S2).

As a result, traditional linear deconvolution methods often struggle to model the biological variability required for the linearity assumption to hold across diverse samples and contexts.

Even when the linearity assumption is valid, several practical challenges remain regarding the creation of the signature matrix. Collinearity in the signature matrix can severely hinder the stability of linear regression models, making the system ill-conditioned. To mitigate this, one must carefully select genes that are both informative and non-redundant, often removing noisy or highly correlated genes. These constraints may lead to the exclusion of biologically relevant genes (Wang, 2019). Furthermore, many linear methods implicitly favor genes with higher mean expression, potentially overlooking genes that are highly discriminatory between cell types but expressed at lower levels (Nguyen, 2024; Tsoucas, 2019). This can limit the resolution and accuracy of the deconvolution. As a result, while constructing a robust and high-quality signature matrix is common practice, it may not always be sufficient due to these inherent limitations.

### 1.2. A latent space approach to deconvolution

Our method directly addresses the core limitations of traditional deconvolution by embedding gene expression data into a latent space designed to approximate ideal deconvolution properties:

- Gaussian-like latent structure.
- Linearity between cell-type specific profiles and mixtures of cells.
- Non-linear mappings that capture subtle, non-linear gene - cell-type dependencies across diverse contexts and retain informative signals from rare or noisy cell types.

To achieve this, we leverage variational autoencoders (VAEs) (Kingma, 2013) to learn a latent representation of gene expression that satisfies both statistical assumptions and biological constraints. Building on single-cell deep learning frameworks such as scVI, we introduce a critical modification to enforce linearity in the latent space - a property not naturally satisfied by standard VAEs.

The key novelty in our approach is the incorporation of a mixup penalty (Beckham, 2019; Berthelot & Raffel, 2018; Khan, 2022; Verma & Lamb, 2018) in the evidence lower bound (ELBO) objective that explicitly optimizes the VAE to jointly model single-cell profiles and their mixtures (pseudobulks) in the latent space. This results in:

- A non-linear encoder that maps both individual cells and mixtures (e.g., bulk RNA-seq, spatial transcriptomics spots) into a shared Gaussian latent space.
- Linearly additive structure in the latent space, where mixture representations reflect their underlying cell type proportions.
- A shared decoder that accurately reconstructs both individual and mixed expression profiles, while modeling their distinct statistical characteristics - such as the zero-inflation commonly observed in single-cell data but not in mixtures.

By training on both single-cell data and systematically generated pseudobulk mixtures, our model learns to “fill the gaps” in the latent space, ensuring that mixtures fall precisely where linear interpolation would predict. This contrasts with standard VAE approaches that may learn efficient representations of individual cells but provide no guarantees about the behavior of mixtures in the latent space.

Our framework thus creates a unified representation where the same latent variables can represent both individual cells and mixtures of cells while preserving the critical property of linear additivity. This enables robust and accurate deconvolution across varying levels of cell type heterogeneity and proportion.

## 2. The MixupVI probabilistic model

For a given single-cell *i*, scRNA-seq experiments output a set of read counts representing the gene expression levels across *G* genes. Let’s denote the gene expression profile of a single-cell as *x*_*i*_ ∈ ℕ^*G*^. Similarly, for a bulk sample *b*, bulk RNA-seq provides a gene expression profile *y*_*b*_ ∈ ℕ^*G*^. We assume each cell *i* is annotated with a cell type label *c*_*i*_ from a set of *C* distinct cell types.

The key insight of our approach is that bulk samples can be represented as a mixture of cells from different cell types, and this mixture relationship should be preserved in the latent space. To achieve this, we design a latent space with an additive property: the representation of the sum of a set of single-cells is approximately the sum of their individual representations.

### 2.1. Generative model for single-cell transcriptomics

Building upon the scVI framework (Gayoso et al., 2022), we first define a generative model for single-cell data:

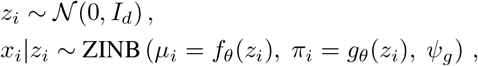

where *z*_*i*_ ∈ ℝ^*d*^ is a *d*-dimensional latent variable representing the cell state, *I*_*d*_ is the *d*-dimensional identity matrix, ZINB denotes the zero-inflated negative binomial distribution, and *f*_*θ*_, *g*_*θ*_ are neural networks parameterized by *θ*, which map the latent variable to the expected gene counts *µ*_*i*_ and the expected dropout rate *π*_*i*_, respectively. *ψ*_*g*_ is a gene-specific dispersion parameter, learned directly for each gene and shared across cells. This formulation captures the over-dispersed and zero-inflated nature of scRNA-seq data.

As with standard VAEs, the marginal likelihood *p*(*x*_*i*_) is intractable, meaning we cannot directly compute the posterior distribution *p*_*θ*_(*z*_*i*_|*x*_*i*_). Instead, we learn an approximate posterior *q*_*φ*_(*z*_*i*|_*x*_*i*_) using variational inference (Kingma & Welling, 2019), where *φ* denotes the parameters of an inference network. We then optimize the evidence lower bound (ELBO) of log *p*_*θ*_(*x*_1:*N*_) with respect to the variational parameters *φ* and the generative model parameters *θ*.

### 2.2. Mixup regularization: enforcing linearity in the latent space

The key novelty in MixupVI is the introduction of a linearity constraint in the latent space, called the mixup loss (Carratino, 2020).

Given a batch of *N* cells and *C* cell types, let’s denote:

- *x*_*i*_ as the *i*-th cell in the batch.
- *z*_*i*_ as the latent representation of cell *x*_*i*_.
- *n*_*cpp*_ defines the number of single-cells to aggregate when generating a single pseudobulk sample. By adjusting this hyperparameter, one can flexibly simulate different experimental resolutions: larger values mimic bulk RNA-seq conditions, while smaller values emulate spot-based spatial transcriptomics where each spot contains fewer cells. This allows for benchmarking or modeling across a spectrum of transcriptomic data types.
- *β* = (*β*_1_, …, *β*_*C*_) as the proportions sampled from a Dirichlet distribution over the *C* − 1 dimensional simplex (i.e., non-negative values that sum to one). This sampling step enables the over-representation of rare cell types, which are often under-represented in classical deconvolution methods due to their low abundance, resulting in insufficient signal. The parametrization of this Dirichlet distribution is detailed in Supplementary Section B.
- ℐ as the set of sampled cell indices for creating a pseudobulk, determined by the proportions *β* and the number of cells per pseudobulk *n*_*cpp*_.

As depicted in Figure 1, the process involves:

**Figure 1.**
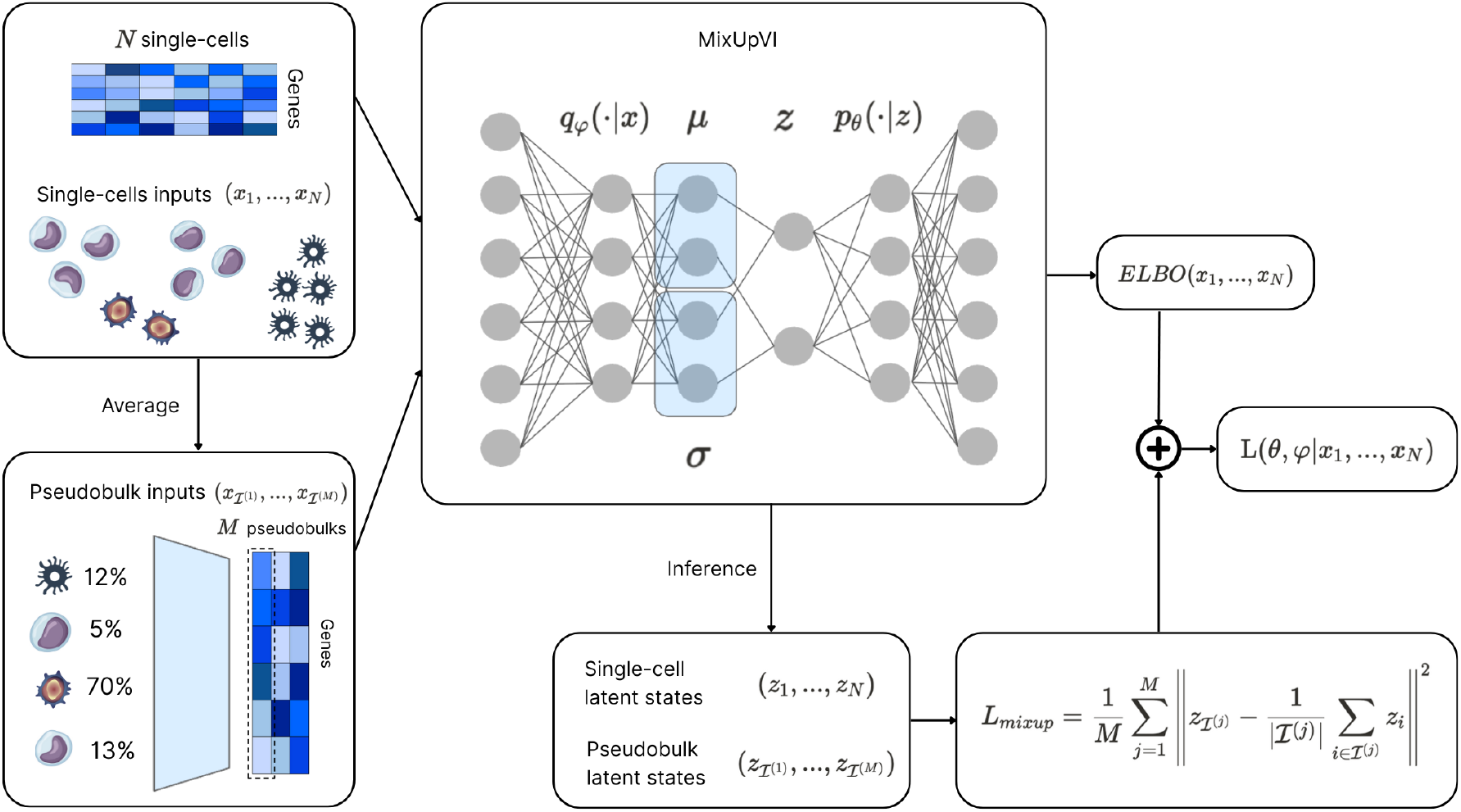
Schematic overview for the MixupVI forward process. Pseudobulks are first created from the batch of single-cells by sampling proportions independently from a Dirichlet distribution and sampling cell indices from the batch to match these proportions. Both single-cells and pseudobulks are then passed to the VAE. Their latent representations are sampled from the encoder to compute the mixup loss. Finally, the mixup loss is added to the ELBO.

- Sampling proportions *β* from a Dirichlet distribution.
- Sampling cell indices ℐ from the batch to match the determined proportions given by *β* and *n*_*cpp*_ with |ℐ| = *n*_*cpp*_.
- Creating a pseudobulk *x*_ℐ_ as a linear combination of the cells, by averaging the sampled cells: 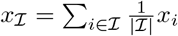.
- Sampling the latent representations *z*_*i*_ ∼ *q*_*φ*_(· | *x*_*i*_) for each cell and *z*_*ℐ*_ ∼ *q*_*φ*_(·|*x*_*ℐ*_) for the pseudobulk, using the encoder of the model.
- We encourage the latent representation of this linear combination to be close to the linear combination of the individual latent representations with a L2 norm. This L2 regularization is particularly meaningful in the latent space, given that the data is modeled as following a Gaussian distribution. Minimizing this penalty is thus equivalent to maximizing the likelihood estimate of the latent variables.

#### Definition 2.1.

For one given collection of cell indices ℐ, the mixup loss can be formulated as follows:

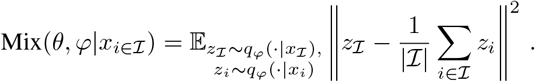

#### Definition 2.2.

To minimize the expected risk over the pseudobulk distribution, we define the mixup loss for the full batch of cells with:

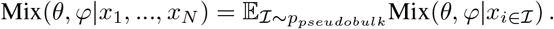

with *p*_*pseudobulk*_ the distribution created from the multi-step stochastic process described above. Collections of indices are drawn according to this distribution. We approximate the resulting expectation using Monte Carlo sampling:

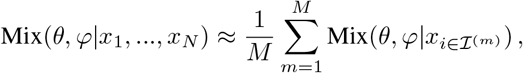

where ℐ^(1)^, …, ℐ^(*M*)^ are sampled i.i.d. following *p*_*pseudobulk*_. This mixup loss term is added as a regularization term to the ELBO:

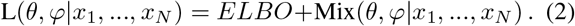

### 2.3. Cell type proportions inference

Given a pseudobulk sample *y*_*b*_, we model it as a mixture of cell types with proportions 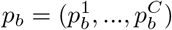:

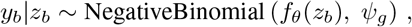

where *z*_*b*_ the inferred latent state for sample *b*.

As depicted in Figure 2, we encode the bulk sample into the latent space *z*_*b*_∼ *q*_*φ*_(·|*y*_*b*_). We then estimate the cell type proportions *p*_*b*_ ∈ ℝ^*c*^ by solving the following constrained linear system (and then dividing the estimated coefficients by their sum to get proportions summing up to 1):

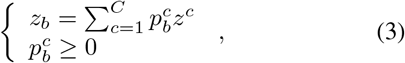

where *z*^*c*^ is the average latent representation for cell type *c* obtained from the single-cell data. This average latent representation serves as the equivalent of a signature matrix in latent space — which we refer to as the latent signature matrix. Unlike classical approaches, it bypasses the need to manually curate a signature matrix, a process that is prone to issues like collinearity and bias toward highly expressed genes.

**Figure 2.**
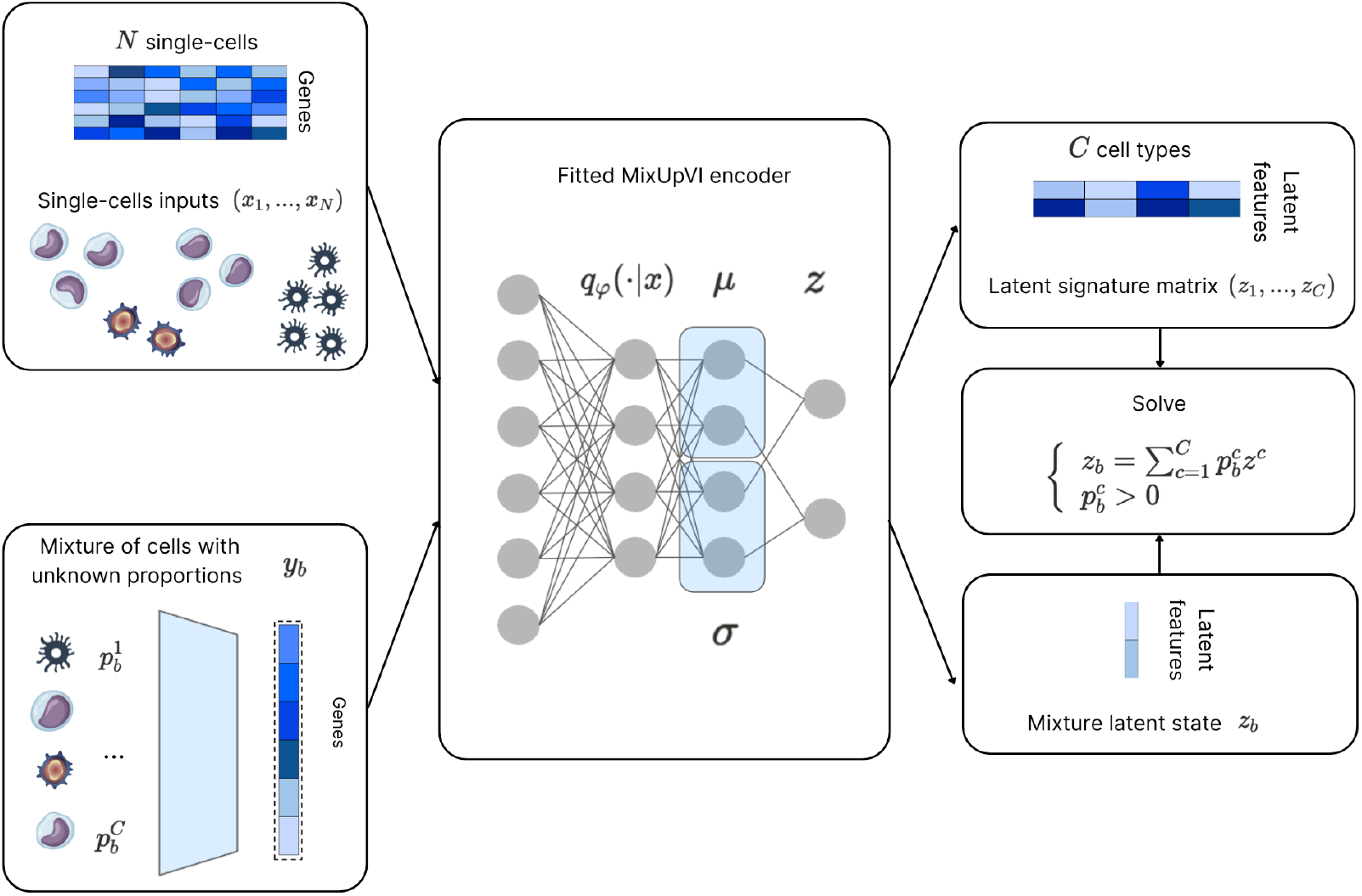
Schematic overview for the cell type proportions inference with MixupVI. The single-cell training dataset is passed to MixupVI. Aggregating the latent representation of the individual cells per cell type allows the creation of the latent signature matrix. The evaluation pseudobulk is passed to MixupVI to infer its latent representation. Finally, the unknown proportions of the pseudobulk sample are inferred with the non-negative least squares approach.

## 3. Results

### 3.1. Datasets and pseudobulk simulation

Our analysis was conducted using the Cross-tissue immune (CTI) single-cell dataset (Conde et al., 2022). This analysis was performed on the immune compartment of 16 tissues from 12 adult donors by single-cell RNA sequencing, creating a dataset of approximately 360,000 cells. For each granularity, we divided the dataset between train and test (50/50 split), using the train set to:

- Build signature matrices (Hao, 2021). We performed differential expression analysis (DEA) between cell types in the single-cell RNA-seq dataset using a non-parametric Wilcoxon rank-sum test on a gene-by-gene basis. Genes were tested only if expressed in at least 1% of cells in either group (cell type of interest vs rest). Differentially expressed genes were defined as those with an adjusted p-value below 0.05 and an absolute log fold change greater than 0.25. Then, minimization of the condition number was conducted using the kappa function. Values reported in the signature matrix correspond to mean gene expression of the significantly differentially expressed genes.
- Fit deep learning algorithms. MixupVI as well as other deep baseline methods (Chen & Wang, 2022; Menden, 2020) included in the benchmark are trained on this set of single-cells.

To ensure robust evaluation, we use the test set to create synthetic pseudobulk datasets with two levels of difficulty defined by cell type granularity (5 and 9 cell types). Pseudobulk samples with known cell type proportions were generated by aggregating single-cells from the test set of the CTI atlas, using proportions given by sampling from a Dirichlet distribution as explained in Supplementary Section B. The breakdown of the cell types for each granularity is shown in Figure 3. We design two types of benchmarks. The first keeps the number of cells per pseudobulk fixed at 100, intentionally inducing under-representation of certain cell types. This choice is motivated by our observation that using larger pseudobulks (e.g., 200–300 cells or more) results in more stable cell type proportions, making the deconvolution task less challenging. The second benchmark varies the number of cells per pseudobulk from 10 to 1000, enabling a comparison of methods across a range of resolution levels.

**Figure 3.**
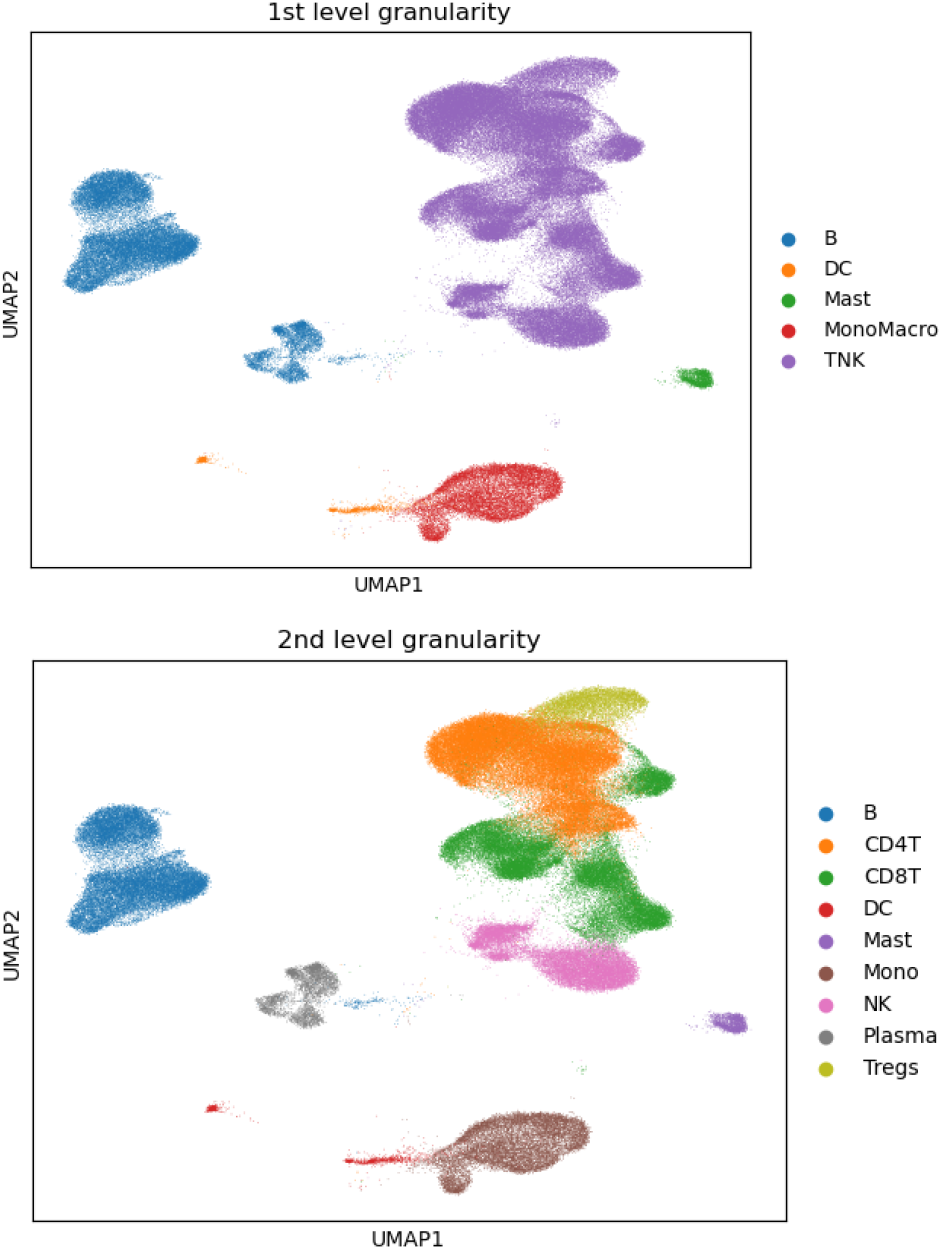
UMAP plot of the embedded single-cell gene expression profiles with scVI. For the 1st granularity level, the cell types grouped are B cells, Dendritic cells, Mast cells, Monocytes & Macrophages, T & Natural Killer cells. For the 2nd granularity level, the cell types grouped are B cells, Plasma cells, Dendritic cells, Mast cells, Monocytes, CD8 T cells, CD4 T cells, T regulatory cels, Natural Killer cells.

### 3.2. PyDeconv: a library for deconvolution methods benchmarking

Our comparative analysis included linear methods: Ordinary Least Squares (OLS), Non-Negative Least Squares (NNLS), Robust Linear Regression (RLR), Nu-Support Vector Regression (NuSVR) which is the core algorithm of CIBERSORT (Newman, 2015), Weighted Non-Negative Least Squares (WNNLS) and Dampened Weighted Least Squares (DWLS) (Tsoucas, 2019). We also included two deep learning methods: TAPE (Chen & Wang, 2022) and Scaden (Menden, 2020). More information about the methods can be found in Supplementary Section C.

Current deconvolution methods are implemented in various programming languages, leading to fragmented workflows. While unified benchmarking is possible using solutions like multi-container Docker pipelines (Nguyen, 2024), such setups can add overhead and require additional maintenance. To streamline this process, we introduce PyDeconv, a Python package that reimplements a set of representative methods along with a benchmarking pipeline using various pseudobulk simulations. The aim of the package is to provide unified support for all major deconvolution techniques found in the literature, thereby simplifying benchmarking efforts and enhancing method robustness through an open-source and standardized framework.

### 3.3. Linearity constraint validation

We first evaluated the effectiveness of our linearity constraint in the latent space. Using the CTI dataset, we trained our model and compared it to the standard scVI model (without the mixup linearity constraint). For validation, we generated synthetic bulk samples (pseudobulks) by combining single-cells from a held out set, in proportions sampled from a Dirichlet distribution and assessed how accurately the models preserved these proportions in the latent space.

We monitored the Evidence Lower Bound (ELBO) on a held out set to ensure that our linearity constraint did not compromise the model’s ability to reconstruct gene expression profiles. To assess the impact of the mixup loss, we measured the Pearson correlation between pseudobulks of the encoded single-cells and the encodings of the pseudobulks created in the input space, that we call mixup correlation *ρ* (Supplementary Section D). Additionally, we computed the Pearson correlation between the estimated cell-type proportions within the latent space and the ground truth proportions for pseudobulk simulated from the held-out set, that we call deconvolution correlation *d* (Supplementary Section D).

Both models exhibited comparable ELBO convergence patterns during training (Supplementary Figure S3). The final values of key metrics on the held out set are reported in Table 1. During training, scVI showed a decrease in mixup correlation *ρ* resulting in an average correlation of 0.32 on the held out set. In contrast, MixupVI maintained a high correlation during training resulting in an average of 0.94 on the held out set.

**Table 1.** Monitoring of validation metrics: ELBO loss, Pearson correlation between encoded pseudobulks in the latent space, Pearson correlation between ground truth and estimated proportions in the latent space at the end of the training.

| METRIC | scVI | MIXUPVI |
| --- | --- | --- |
| ELBO LOSS | 641.16 | 641.37 |
| MIXUP CORRELATION $\rho$ | 0.32 | 0.94 |
| DECONVOLUTION CORRELATION $d$ | 0.45 | 0.88 |

Furthermore, these correlations in the latent space translate to deconvolution accuracy of synthetic pseudobulks generated from the held out set. scVI demonstrated poor deconvolution performance throughout training, while MixupVI improved its deconvolution capabilities: by the end of training, MixupVI achieved a deconvolution correlation *d* close to 0.9, twice as much as scVI.

These results indicate that MixupVI maintains reconstruction quality while gaining the critical linearity property necessary for accurate deconvolution.

### 3.4. Comprehensive benchmarking of deconvolution methods

The metrics presented in this part are the deconvolution correlation *d* and Mean-Squared Error (deconvolution MSE) between the estimated cell-type proportions of each deconvolution method and the ground truth proportions for pseudobulk simulated from the held-out set (Supplementary Section D).

#### First-level granularity (5 cell types)

At the coarse granularity level distinguishing major immune cell compartments, MixupVI demonstrated competitive performance with other best performing methods (RLR, WNNLS, Scaden) with a median Pearson correlation of 0.973 (Figure 4.a.). Naive linear methods such as NNLS and OLS, which do not incorporate weighting, tend to underperform. This could be due to collinearity issues in the signature matrix as well as the limitations discussed in the introduction: broad cell type definitions often exhibit high intra-cell type variability, causing the estimated cell type profiles to deviate from those in the signature matrix. Notably, these methods show high Pearson correlation variability and occasionally produced negative correlation values, highlighting their instability for this task.

**Figure 4.**
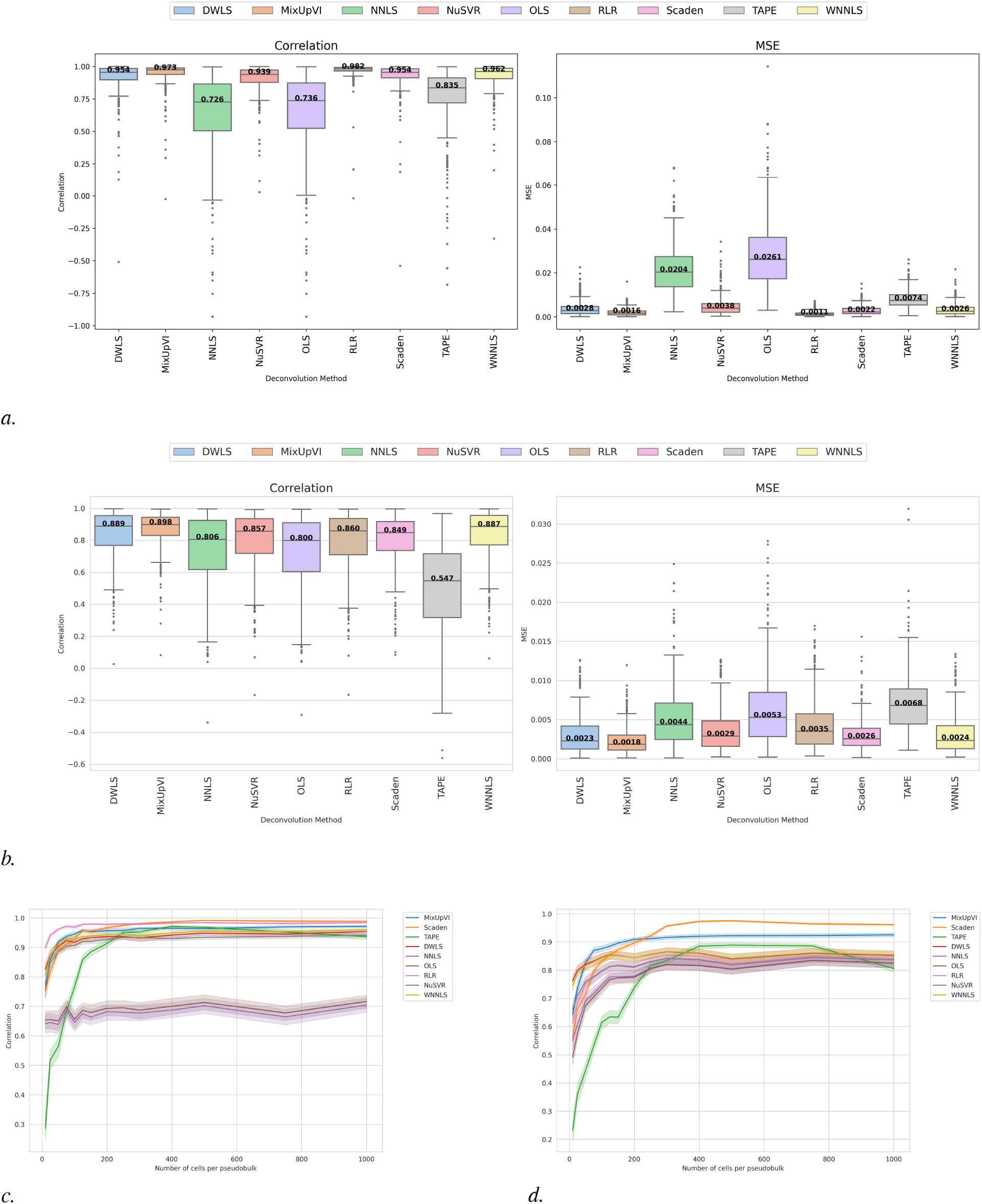
Deconvolution results. *a*. 1st granularity results. Pearson and MSE of predicted vs groundtruth proportions (400 pseudobulk samples / 100 cells per pseudobulk). *b*. Same metrics for 2nd granularity. *c*. Trend of the deconvolution results with regards to the number of cells samples for each pseudobulk simulation for the 1st granularity. The y-axis shows the average Pearson correlation and on the x-axis number of cells per pseudobulk. *d*. Same metrics for 2nd granularity.

RLR and MixupVI also achieved the lowest MSE at 0.0011 and 0.0016 respectively, outperforming all other competing methods.

#### Second-level granularity (9 cell types)

When the deconvolution task difficulty was increased to distinguish finer cell subtypes (including separation of CD4^+^ T cells, CD8^+^ T cells, T regulatory cells, and Natural Killer cells), MixupVI maintained robust performance and low variability, with a median correlation of 0.898 (Figure 4.b.). This represented a marked improvement of at least 1 point over all competing methods.

The MSE results further confirmed MixupVI’s advantage, with a median error of 0.0018 compared to 0.0026 for Scaden and 0.0023 for DWLS. The performance gap between MixupVI and other methods widened at this higher granularity level, demonstrating our method’s particular advantage in more challenging deconvolution scenarios.

#### Varying number of cells per pseudobulk

In addition to our primary benchmark results, we investigated how deconvolution performance varies with the number of cells in each pseudobulk sample. This analysis provides important insights into method robustness across varying input data depths.

At the first-level granularity (Figure 4.c.), besides OLS, NNLS and TAPE, most deconvolution methods are competitive, showing similar trends with increasing performance across sample sizes, until it plateaus around 100 cells. In higher sample size regime (*>* 250) Scaden outperforms other competing methods, which is likely due to Scaden’s higher default sample size of 500 used in the simulations during training. In contrast, NNLS and OLS performs poorly across sample sizes, highlighting the limitations of classical methods when cell type definitions are overly broad, as averaging across heterogeneous populations can obscure important expression differences and degrade deconvolution accuracy.

At the second level of granularity (Figure 4.d.), MixupVI is outperformed by weighted linear methods when the number of cells is very low (*<* 25). However, for sample sizes between 25 and 200 cells, MixupVI consistently outperforms all other methods, highlighting its robustness to input variability, data sparsity, and the under-representation of rare cell types. Similarly to the 1st granularity experiments, we also observe that the the performance of Scaden steadily increases until outperforming other methods in the high cell-count regime (*>* 250). These findings underscore MixupVI’s robustness to input data diversity and sparsity — an important advantage in real-world settings where high-depth bulk RNA-seq data may be difficult to obtain or where only limited samples are available. This robustness also suggests a potential benefit for spatial transcriptomics deconvolution, where each spot captures a limited number of cells. The consistent performance advantage across varying granularities and sample sizes demonstrates the broad applicability of our approach to diverse deconvolution scenarios.

## 4. Discussion

### 4.1. Impact of the latent size

MixupVI is designed to be easily adaptable to new datasets, data modalities and levels of granularity. In Supplementary Section E, detailed explanations of every benchmark, training procedure, and model hyperparameters are provided. However, among the various model hyperparameters, the latent space size is particularly crucial, requiring careful tuning.

As shown in (1), the signature matrix used with Non-Negative Least Squares (NNLS) is of size (*n*_*genes*_, *n*_*cell types*_). Similarly, in the case of MixupVI, the latent signature matrix used with NNLS is of size (*n*_*latent size*_, *n*_*cell types*_). Thus, to avoid high-dimensionality issues, the latent space size should be at least higher than the number of cell types. Plus, it should be chosen to remove noise or collinearity of gene expressions that could impede fitting. Classical methods achieve this by carefully constructing the signature matrix. MixupVI, on the other hand, embeds gene expressions in the latent space to reduce noise and collinearity. Finally, the latent space should be sufficiently large to allow flexible shaping of representations through complex non-linear mappings, as the mixup loss introduces structural constraints that limit representational freedom.

The impact of the latent size is apparent in Supplementary Figure S4, showing deconvolution correlation *d* and deconvolution MSE during training, on synthetic pseudobulks generated from the validation set. On the 9 cell types granularity, the performance improves progressively with a larger dimensionality for both metrics: a dimensionality of 30 yields improved performance compared to 20 and 10. We also notice that performance plateaus with this parameter - for instance when the latent space is increased to 100, the MSE plateaus, and even worsens for the Pearson correlation.

### 4.2. Conclusion

We have presented MixupVI, a deep generative model that learns probabilistic representations of single-cell RNA-seq data with a linearity constraint enabling accurate deconvolution of mixtures of cell types. By enforcing an additive property in the latent space, our model effectively bridges the gap between the expressivity of deep generative models and the linearity requirements of deconvolution.

MixupVI offers several key advantages over existing deconvolution methods. First, it constructs a latent Gaussian space in which samples are naturally expressed as mixtures of constituent cell types, aligning with the assumptions underlying deconvolution. Second, it models biological variability within cell types via its latent representation, enhancing accuracy - especially when distinguishing closely related types. Third, its probabilistic framework yields uncertainty estimates for both cell type proportions and reconstructed gene expression profiles, enabling more informed interpretation. Finally, it eliminates the need for manually curated reference profiles by learning a latent representation directly from the data, avoiding common pitfalls of signature matrix construction. Given its reliance on a latent space, however, the dimensionality of this space becomes an important design choice that should be tailored to each deconvolution task.

Future work will aim to extend MixupVI to accommodate more complex and realistic settings. Modeling pseudobulks with a small number of cells would enable its application to spot-based spatial transcriptomics, while larger pseudobulks make it suitable for bulk RNA-seq paired with flow cytometry. However, key technological differences between bulk, single-cell, and spatial transcriptomics must be carefully addressed to adapt MixupVI for real-world multi-omics integration and ensure meaningful cross-modality alignment.

## Software and Data

All the referenced methods, the pseudobulk simulations and the benchmarking code were implemented in the python library PyDeconv github.com/owkin/pydeconv.

The Cross-Tissue Immune atlas was dowloaded from cellx-gene.

## Acknowledgements

We would like to express our deep gratitude to Sabrina Carpentier for her unwavering support, optimistic guidance, and valuable insights throughout the project, particularly regarding data generation processes. We also thank Ulysse Marteau for his initial mathematical formalization of the deconvolution framework and the mixup approach. Finally, we warmly thank Alessandro Pranzo, who has now enthusiastically joined our team to help enhance the model and pipelines.

## Impact Statement

This paper presents work whose goal is to advance the fields of Machine Learning and Biology. There are many potential societal consequences of our work, none which we feel must be specifically highlighted here.

## A. Supplementary figures

**Figure S1.**
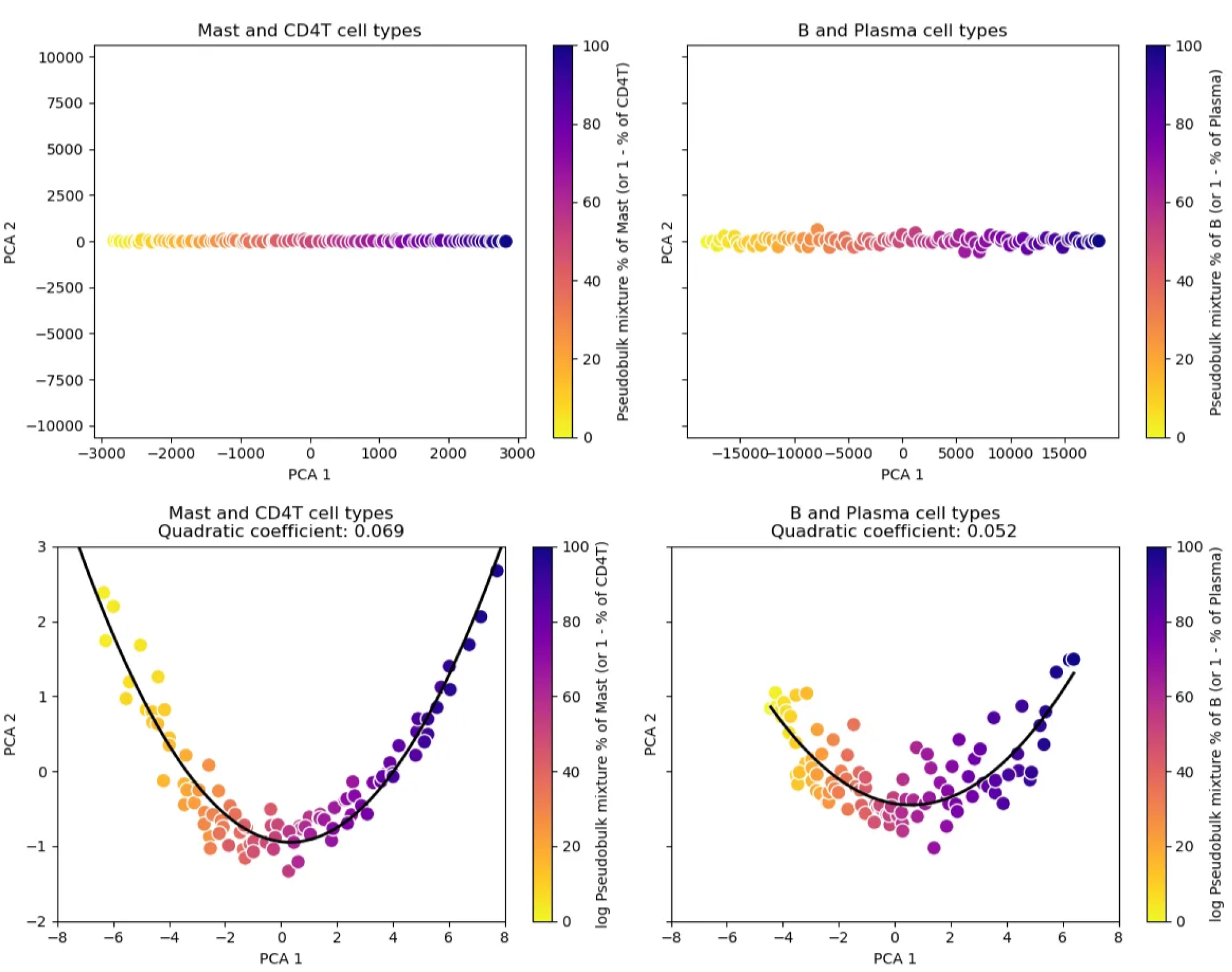
PCA of mixed pseudobulks. With the Cross-tissue immune single-cell atlas (section 3.1), we construct pseudobulks composed of varying proportions of cell types. The interpolation between mixtures is linear when the gene expression is not log-transformed (above figure), while the interpolation between mixtures shows a curved trajectory when the gene expression is log-transformed (below figure). The effect is more pronounced when combining dissimilar cell types, such as Mast and CD4T cells, compared to more similar ones like B and Plasma cells.

**Figure S2.**
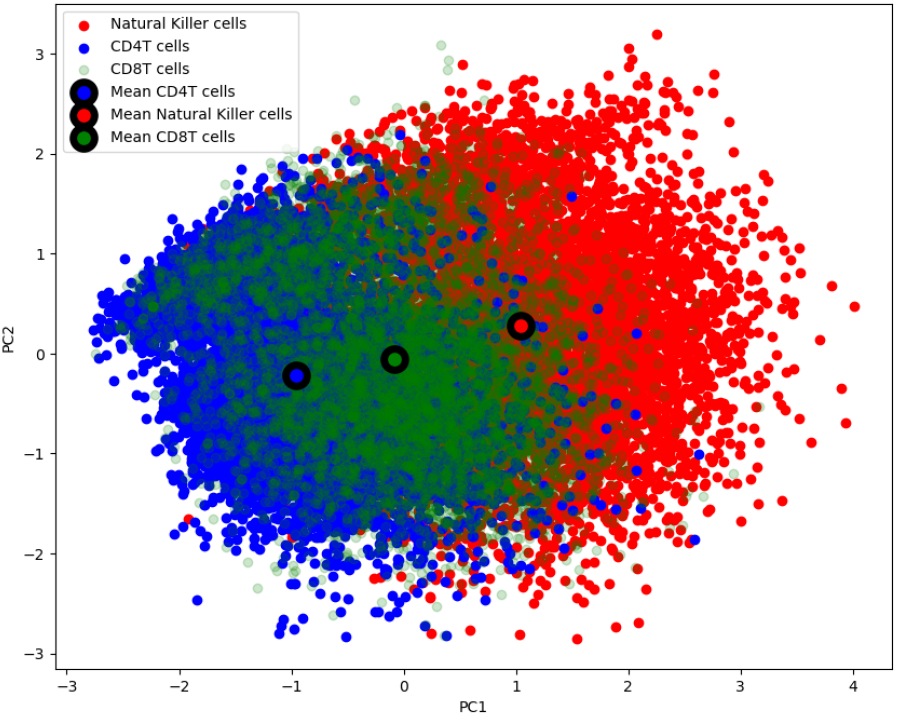
Using the Cross-tissue immune single-cell atlas (Section 3.1), we analyze the variability of lymphoid cells gene expression. Due to shared functions and markers, CD8^+^ T cells were grouped with both CD4^+^ T (Wang, 2008) and NK cells (Rosenberg, 2017) in the broader granularity. While CD4^+^ T and NK cells are mainly separated along PC1, they are both mixed with CD8^+^ T cells. This highlights underlying similarities between CD4^+^ T and NK cells, but also sufficient differences that may introduce unwanted variability in coarse cell type definitions.

**Figure S3.**
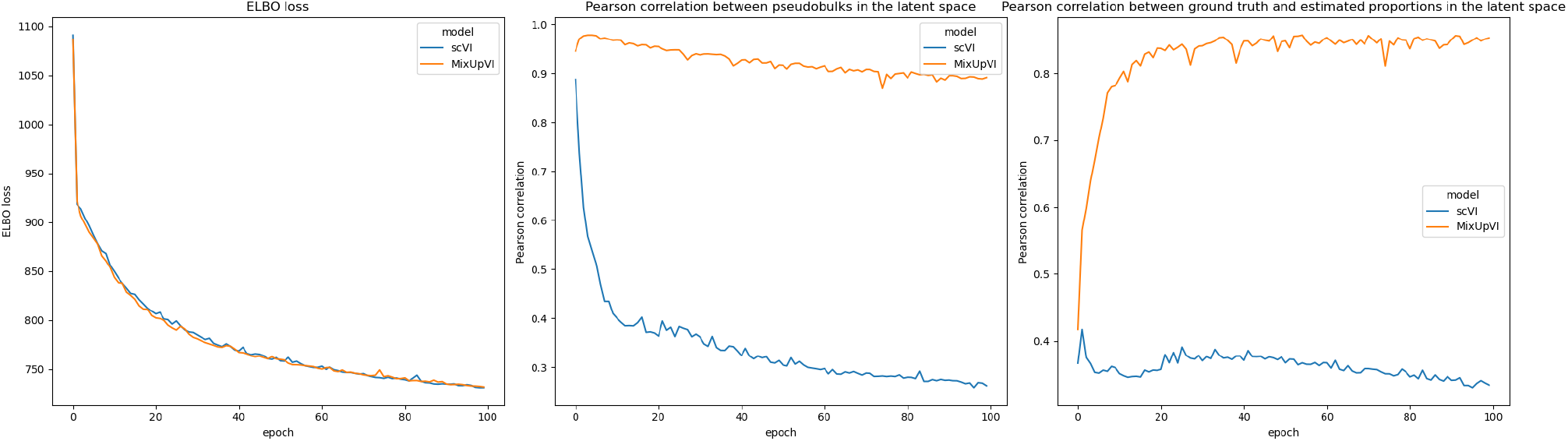
Monitoring of the ELBO loss (left plot), Pearson correlation between encoded pseudobulks in the latent space (middle plot), Pearson correlation between ground truth and estimated proportions in the latent space (right plot) during training.

**Figure S4.**
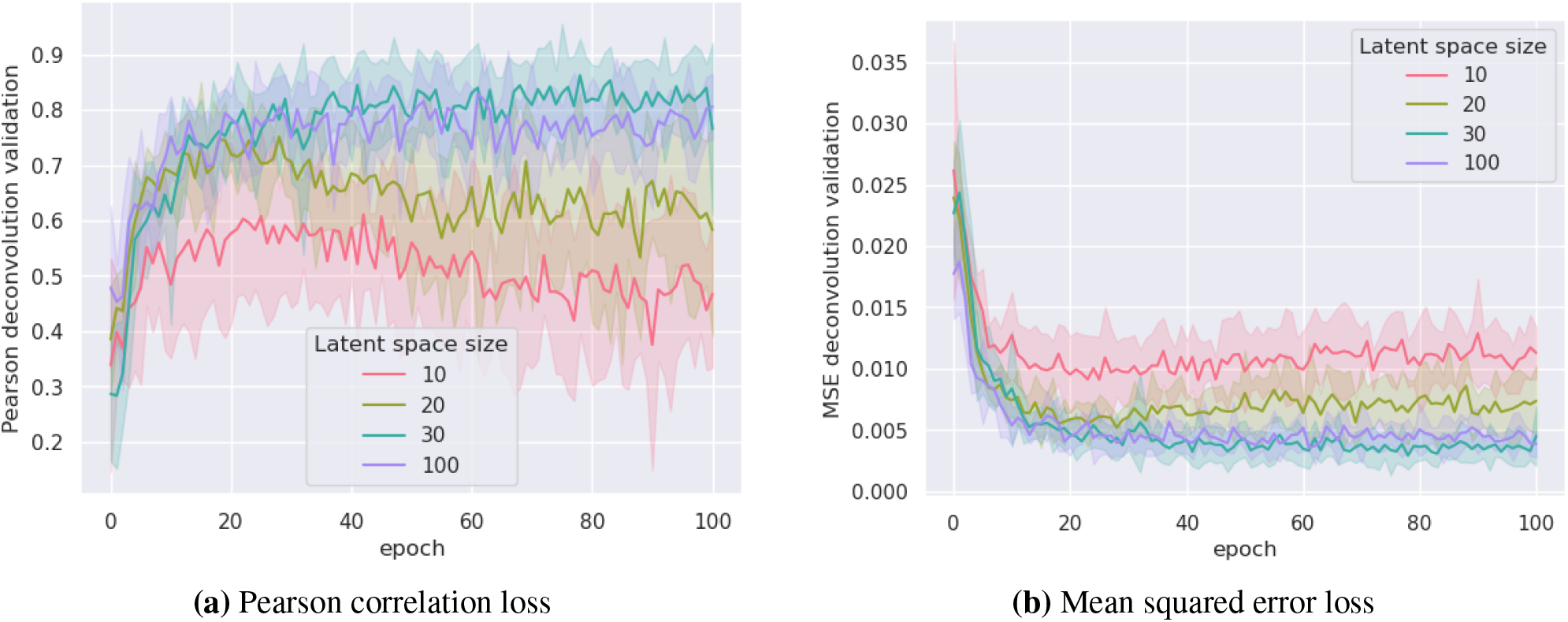
Example of latent space size tuning for the 9 cell types granularity. The deconvolution correlation *d* (a) and deconvolution MSE (b) are computed at every batch of every epoch on the validation set. The error band is the standard deviation across 5 different seeds. Intermediate latent sizes (between 30 to 100) are not shown for clarity.

## B. Dirichlet sampling

As explained in Section 2.2, we sample the pseudobulk cell type proportions from a Dirichlet distribution. This is a useful sampling in the case of pseudobulk creation because it sums proportions to 1 and allows for over-representation of rare cell types.

This Dirichlet distribution is the posterior of a Dirichlet-Multinomial conjugate model, assuming a non-informative prior and a multinomial likelihood based on the observed cell counts. More precisely:

- We assume that (*p*_1_, …, *p*_*C*_) to be the vector of multinomial parameters, thus the probabilities associated to every cell type.
- We assume that this vector follows a Dirichlet prior to observing the batch of single-cells. More formally: (*p*_1_, …, *p*_*C*_) ∼ *Dirichlet*(*α*_1_, …, *α*_*C*_).
- In this benchmark, we chose a non-informative prior distribution: *α*_1_ = … = *α*_*C*_ = 1. In that special case, the Dirichlet distribution is actually a Uniform distribution. It is also possible to set this prior to match the belief one has over the cell type proportions of the given tissue from which the single-cell data is drawn.
- Then, a batch of single-cell is drawn. So are the cell type proportions of this given batch: *x*_1_, …, *x*_*C*_. Given these observed cell type proportions, using the conjugate property of the Dirichlet and Multinomial distributions, we can easily compute the posterior distribution of the vector of multinomial parameters:

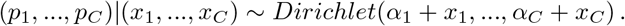

## C. Related work

As detailed in Section 3.2, we benchmarked MixupVI against a broad range of established deconvolution methods (Cobos, 2020), all provided as Python implementations in our open-source library PyDeconv. This includes both classical regression-based techniques and deep learning models. Several of these methods were originally implemented in R, but have been re-implemented with Python to ensure consistency and accessibility. Below is an overview of each method:

- Ordinary Least Squares (OLS) A standard linear regression technique that estimates cell type proportions by minimizing the squared error between the observed bulk expression and the weighted combination of reference cell-type profiles. While simple and fast, OLS does not constrain predictions to be non-negative or sum to one, which can lead to biologically implausible results.
- Non-Negative Least Squares (NNLS) An extension of linear regression that constrains all estimated cell type proportions to be non-negative. This makes NNLS more realistic for compositional data like cell mixtures and has been widely used as a baseline in deconvolution studies.
- Robust Linear Regression (RLR) This method modifies standard regression by reducing the influence of outliers or highly variable genes. It improves stability in the presence of noise or batch effects, which are common in real-world transcriptomic datasets.
- Weighted Non-Negative Least Squares (WNNLS) W-NNLS improves upon NNLS by incorporating gene-specific weights based on their variance across cell types. This prioritizes genes with higher discriminatory power, enhancing the robustness of proportion estimates.
- Dampened Weighted Least Squares (DWLS) DWLS extends WNNLS by further iteratively adjusting the contribution of each gene through dampening weights, reducing the impact of genes with extreme variance. This method is particularly effective for detecting rare cell populations and managing noise in lowly expressed genes.
- Nu-Support Vector Regression (NuSVR) This model is based on a linear kernel model using an extra regularization parameters nu which simultaneously controls the number of support vectors and the upper bound on the fraction of training errors. Compared to OLS, the SVR model is robust against noise, can automatically select important genes from the signature matrix, and can account for multicollinearity between cell types.

In addition to classical regression-based methods, we benchmarked two deep learning approaches that model complex non-linear relationships in gene expression data:

- Tissue-Adaptive autoEncoder (TAPE) TAPE leverages an autoencoder-based architecture trained on simulated bulk RNA-seq data to learn latent representations that capture cell-type specific expression patterns. The training phase uses deep neural networks trained on pseudobulk datasets derived from single-cell RNA-seq data. During inference, TAPE incorporates an adaptive self-supervised step to perform domain adaptation, taking advantage of the autoencoder reconstruction capabilities to better generalize to real bulk samples.
- Scaden Scaden employs an ensemble of deep neural networks trained on simulated pseudobulk datasets derived from single-cell RNA-seq data. As with most neural network approaches, its performance relies on access to a large and diverse dataset in order to generalize well across different tissues.

## D. Metrics

To evaluate the impact of the mixup loss and assess latent deconvolution performance, we compute three metrics: two based on Pearson correlation and one based on mean squared error.

First, we measure the alignment between the encoding of a pseudobulk and the average of its constituent single-cell encodings. For a given pseudobulk *x*_ℐ_, let *z*_ℐ_ ∼ *q*_φ_ (·|*x*_ℐ_) denote its latent representation, and let 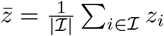, with *z*_*i*_ ∼ *q*_*φ*_(· | *x*_*i*_), be the average of the encoded cells. The mixup correlation for one pseudobulk is then defined as:

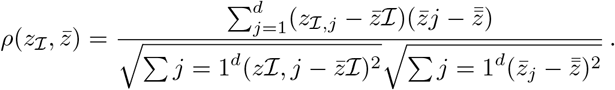

Averaging this over *M* pseudobulks gives the average mixup correlation:

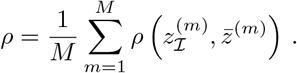

Second, to quantify (latent) deconvolution performance, we compare the predicted proportions 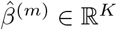 with the true proportions *β*^(*m*)^ ∈ ℝ^*K*^ used to generate pseudobulk *m*. The Pearson correlation for one pseudobulk is:

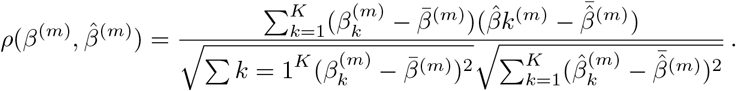

The overall performance across pseudobulks is denoted the deconvolution correlation *d*:

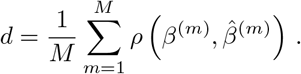

Third, we compute the mean squared error (MSE) between the predicted and true proportions:

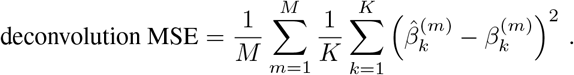

## E. Hyperparameters

Here we describe the parameters of typical benchmarking configs and hyperparemeters used and tuned to train MixupVI.

### Deconvolution benchmark hyperparameters

- Deconvolution method. The deconvolution methods to benchmark in this experiment, among MixupVI, NNLS, scVI, DestVI, TAPE, Scaden, OLS, DWLS, RLR, NuSVR, WNNLS.
- Evaluation datasets. The single-cell datasets to test for deconvolution, used to create pseudobulks. In this benchmark, the test set of the Cross-tissue immune atlas was used.
- Training dataset. The dataset to use to train methods requiring training (e.g. MixupVI). In this benchmark, the train set of the Cross-tissue immune atlas was used.
- Granularities. The granularities to benchmark. In this benchmark, a first granularity with 5 cell types and a second granularity with 9 cell types were tested.
- Signature matrices. The signature matrices to use depending on the granularity. In this benchmark, signature matrices created with the training dataset were used for each granularity.
- Number of evaluation pseudobulks. The number of evaluation pseudobulks simulated for evaluation. In this benchmark, 400 pseudobulks were created.
- Number of cells per evaluation pseudobulk. The number of cells constituting pseudobulk simulations. In this benchmark, a list ranging from 10 cells to 1000 cells was used.

### MixupVI model hyperparameters

#### All scvi-tools models hyperparameters

- Latent size. As described in Section 4.1, this hyperparameter is crucial and should be carefully tuned depending on the granularity. In the 1st granularity, we fixed it to 10, while on the 2nd granularity, it was fixed to 30.
- Number of input genes. The number of input genes to keep in the training datasets for the methods needing training. This parameter is common to all methods requiring training (e.g. MixupVI). It is tunable, as a trade-off needs to be found between high-dimensionality, long training times and preserving cell type markers. In this benchmark, the 2500 most variable genes were kept.
- Maximum number of epochs. The maximum number of epochs to train the model on. In this benchmark, it was fixed to 100 epochs.

#### MixupVI training hyperparameters

- Batch size. This will determine, in the mini-batch gradient optimization, the number of cells in a given batch. By extension, this is the maximum number of cells a training pseudobulk can be composed of. In this benchmark, it was fixed to 2048.
- Train size. For the training dataset described in Section 3.1, this is the proportion kept for purely training, and by extension, it sums up to 1 with the proportion kept for validation at every forward pass. In this benchmark, it was fixed to 0.7.

#### MixupVI model hyperparameters

- Number of pseudobulks. The number of pseudobulks created in an ensembling fashion in a given single-cell batch to compute the mixup loss. The more pseudobulks, the longer a run and the more demanding on GPU memory. In this benchmark, it was fixed to 100.
- Number of cells per pseudobulk. The number of cells constituting the pseudobulks. By adjusting this hyperparameter, one can flexibly simulate different experimental resolutions: larger values mimic bulk RNA-seq conditions, while smaller values emulate spot-based spatial transcriptomics where each spot contains fewer cells, with a better modeling of the variance inside the latent space. In this benchmark, it was fixed to 100.
- Size and number of hidden layers. In the benchmark, one hidden layer was used, of size 512.
- Continuous and categorical covariates. Although theoretically useful to mitigate batch issues, such as center or patient effects (if these effects are present in the training set), they did not improve results in the benchmark, thus they were not used.
- Loss computation. Whether to enforce the mixup loss within the latent space or the reconstructed space (or both). In the benchmark, it was fixed to the latent space.
- Pseudobulk computation. Whether to create the pseudobulks in the input space, or within the latent space, before passing them through the encoder, or decoder. When the loss is computed in the latent space, only the pre-encoded pseudobulks can be computed.
- Signature type. Whether to compute the latent signature matrix used for deconvolution by using pre-encoded purified cell type vectors and passing them through the encoder, or to sample them directly from the latent space. This hyperparameter does not change the optimization of the model, as no deconvolution is done to optimize the model, only to infer from it.
- Mixup penalty. It can either be a L2 loss between the pseudobulk of the sampled encoded cells and the sampled encoded pseudobulk, or the Kullback-Leibler divergence between the distribution of the sampled encoded pseudobulk and the pseudobulk of the sampled encoded cells. In this benchmark, as explained in Section 2.2, the L2 loss was used.
- Gene likelihood. The distribution of the input data, used to compute the reconstruction loss part of the ELBO. In this benchmark, it was fixed to Zero-inflated negative binomial (ZINB) for single-cells and Negative binomial (NB) for pseudobulks.
- Batch normalization. Whether to add batch normalization inside the encoder or/and decoder for every batch of a forward pass. In this benchmark, it was chosen to not use it neither in the encoder or the decoder.

## Notes

https://cellxgene.cziscience.com/collections/62ef75e4-cbea-454e-a0ce-998ec40223d3

### Competing Interest Statement

All authors are employees of Owkin, Inc., New York, NY, USA.

### Summary of Updates

- Proceedings of the ICML conference 2025 added as a footnote. - Two-column format instead of single-column. - Figure 4.b. revised because it did not showcase the right number of cells per pseudobulk. Conclusions and relative order of the methods remain the same but the actual median performance numbers displayed change. - A few wordings.

